# Behavioural changes in female northern bats (*Eptesicus nilssonii*) across gestation and lactation

**DOI:** 10.1101/2025.05.14.653963

**Authors:** Thomas M. Lilley, Mari Aas Fjelldal, Jeroen van der Kooij

## Abstract

Unpredictable environmental conditions affecting prey availability and short summer nights reducing foraging time present nocturnal insectivorous mammals with an energetic dilemma at northerly latitudes during gestation and lactation. This is pronounced for aerial hawking bat species, for which prey activity is highly dependent on ambient temperature. Here, we describe changes in activity of female *Eptesicus nilssonii*, the northernmost breeding bat species, during gestation and lactation over several years at a roost site in southern Norway (60.1°N). Roost exits and returns were recorded using an infrared barrier at the entrance, and parturition dates were registered via a camera set inside the roost. Our results indicate significant temporal changes in body mass and behaviour throughout the breeding season. Lactating bats both setting out to forage earlier than during gestation and returning later. This behaviour is dependent on ambient temperature, with bats extending foraging time at elevated temperatures. These behaviours may occur as a response to the increased energetic needs of lactation, and the effect of increased temperature to reflect increased food availability which allows females to avoid increased predation pressure as a consequence of advanced roost exit or delayed return. The mass of females also decreases during lactation, suggesting high energetic demands during this period. Our study reveals insights into underlying mechanisms in high latitude insectivorous bats that can assist in coping with a short season with fluctuating resources.

## Introduction

Seasonality presents the strongest source of external variation influencing natural systems (Fretwell 1972; Wingfield 2008). It contributes significantly to contemporary biodiversity as well as the evolution of physiological adaptations and behaviours such as hibernation and migration (Varpe 2017). These adaptations have evolved through direct need to facilitate survival and increase fitness in the far north and south, where organisms experience extreme variation in temperature and day length across the annual cycle. Small insectivorous mammals in particular face extreme energetic challenges due to variability in food availability and the limited time window for breeding. These challenges are pronounced in nocturnal mammals with limited foraging opportunities due to reduced night length at a time when insect food becomes available (Fjelldal et al. 2023).

Bats (Chiroptera) are a speciose order of mammals with over 1400 species across all continents apart from Antarctica (Zachos 2020). Although the dietary variability in the order is large, roughly 70% of all bat species are insectivorous (Simmons 2005) and this constitutes the diet of all bat species found at northerly latitudes. When depending on an energy source that varies vastly in availability, survival in temperate regions is enabled by dynamic use of torpor (Dietz and Kalko 2006; Fjelldal et al. 2023), a physiological adaptation that reduces energetic demands through metabolic constraint (Barclay et al. 2001). In spring, while environmental conditions are inhospitable and food availability is highly variable, pregnant females may use daily torpor to help build their depleted reserves after the winter hibernation. However, the use of torpor by pregnant females slows the development of the foetus (Racey and Swift 1981; Dzal and Brigham 2013), which delays parturition. Reproductive females may therefore delay birth of their pups until after foraging conditions have improved (Willis et al. 2006). Bats at northerly latitudes face an imperative to optimize the timing of their pregnancy and lactation with ambient light and weather conditions that maximize foraging time and food availability (Reimer and Barclay 2024).

The northern bat (*Eptesicus nilssonii*) is the northernmost bat species in the world (Suominen et al. 2020), with records of breeding above the arctic circle (Rydell et al. 1994). The species shows tolerance to ambient light (Rydell 1992a; Frafjord 2021), utilises roosts and foraging habitats variably depending on environmental conditions (Vasko et al. 2020; Suominen et al. 2023; Suominen et al. 2024), is a generalist in regard to insect diet (Vesterinen et al. 2018), and makes extensive use of dynamic torpor behaviour across the active season (Fjelldal et al. 2023; Fjelldal et al. 2024; Suominen et al. 2024), facilitating survival at extreme latitudes. *Eptesicus nilssonii* gives birth under higher ambient light conditions in comparison to other records of North-European bat species (Linton and Macdonald 2018), with parturition in mid-summer, close to summer solstice (Rydell 1989; Frafjord 2013; Fjelldal and van der Kooij 2024). Furthermore, pups become volant very rapidly, c. two weeks after birth, and assume adult flight patterns within a week after their first flight (Fjelldal and van der Kooij 2024), adding to their adaptations to the short active seasons.

Due to short mid-summer nights limiting foraging durations for bats at higher latitudes (Frafjord 2013), pregnant and lactating females face a period of restricted foraging opportunities while experiencing elevated energy demands (Kurta et al. 1989; Rydell 1993a). To meet these demands, lactating *E. nilssonii* have been suggested to increase the duration of their foraging activity (Rydell 1993a); however, this increase would indicate that females emerge earlier from their roosts, and/or that they increase the duration or number of foraging flights to return closer to sunrise. Because insect activity at northern latitudes is dependent on ambient temperature and thus rapidly dwindles after sunset, bats emerging earlier benefit from higher insect abundance, but face increased predation risk from diurnal avian predators (Rydell et al. 1996; Speakman et al. 2000). Extending foraging activity towards sunrise instead could minimise risk from diurnal predators but would return less arthropod prey in return for the effort. At northern latitudes with short nights, we expect that the decision of reproductive *E. nilssonii* is likely finetuned towards the cost-benefit scenario with the potential for maximizing individual fitness.

Using data from a single monitored maternity roost of *E. nilssonii* spanning multiple years we investigate temporal changes in roost exits and returns in female bats before and after parturition. We predict bats in the monitored roost to compensate for the higher energetic demands of lactation (Kurta et al. 1989; Mclean and Speakman 1999) during a period when environmental factors are challenging by extending the duration outside the roost, facilitating increased time available for foraging. This can take place by emerging from the roost earlier, by returning to the roost later, or extending at both extremes. Furthermore, we provide additional details on body mass change during gestation and lactation, and how activity of female bats varies in relation to environmental conditions.

## Methods

### Study system

The study utilises a maternity colony (total N_ind_ = 8) of *E. nilssonii* (Fig. 1a) roosting in a bat box, which hangs 1.5 m above ground on a garage-wall in Nittedal, Norway (60.11°N, 10.85°E; Fig. 1b). Data on the colony were collected across the breeding seasons of 2017 to 2023. At this northern latitude, the temporally increasing spring temperature conditions first stabilise around the time of summer solstice, when nights are at their shortest (Fig. 1c), corresponding to the timing of parturition in *E. nilssonii* at this latitude (Fig. 1d).

**Figure 1:**
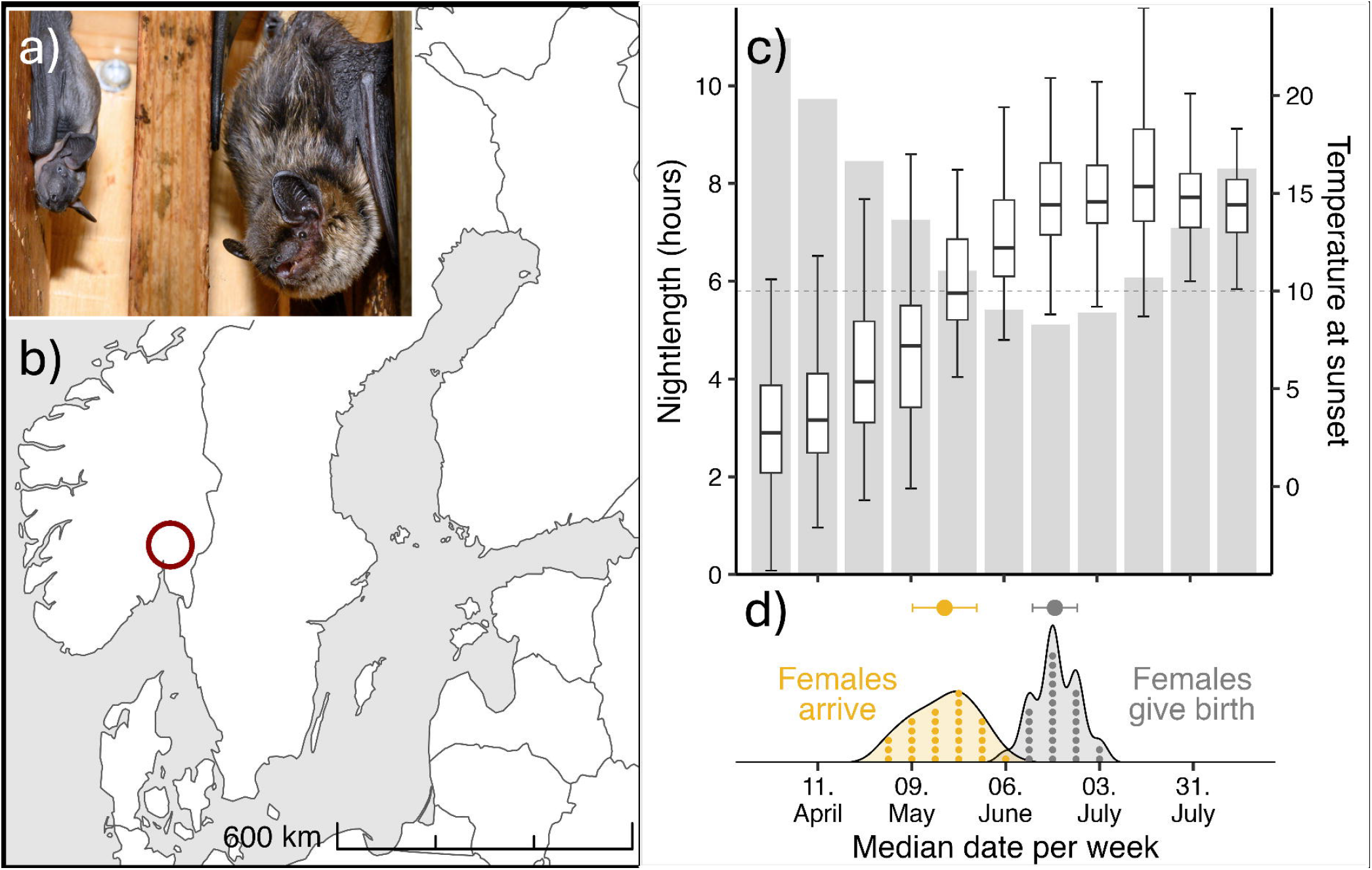
Visual presentation of the study system. **a)** Picture of one of the studied E. nilssonii females with her pup. **b)** Location of study system in Norway, marked with red circle. **c)** Nightlength (shaded bars and left y-axis) and temperature at sunset (boxplots and right y-axis; dashed line indicating 10 degrees at sunset; data from 2017 to 2023) in the study system across the breeding period. **d)** Breeding phenology of E. nilssonii at this latitude, indicating the timing of arrival at the colony (yellow density plot) and timing of parturition (grey density plot). The mean and standard deviation of arrival and parturition are shown above the respective density plots.

Six of the females in the colony had been brought in as abandoned pups and were hand-reared until they could be released. These females returned to the bat box in subsequent years to give birth, and the two last females were daughters born in the box that returned the following summers as part of the maternity colony. Individual identification within and across years was determined based on ring numbers and their placement (two females were ringed as juveniles, one on the left wing and one on the right wing), the individual patterns of collagen-elastin bundles in their wings (Amelon et al. 2017), and on forearm-lengths, given the small number of reproductive females in the box each season (N_ind_ 2017 = 2; N_ind_ 2018 = 5; N_ind_ 2019 = 6; N_ind_ 2020 = 4; N_ind_ 2021 = 4; N_ind_ 2022 = 4; N_ind_ 2023 = 5). One of the ringed females was present in the colony across all study years, while the second ringed female did not return after the breeding season of 2020.

A heating device (135-Watt heater with thermostat) was installed at the top of the box but was turned off on warm days to avoid overheating. We monitored the thermal environment inside the box with iButtons (model DS1923-F5, Dallas Semiconductor Inc., Dallas, TX, USA) during the season of 2020 and confirmed that the thermal gradient was suitable and allowing the bats an opportunity to select a preferred microclimate (mean temperature conditions at the top of the box was recorded to be 29.1°C ± 5.4 *SD*, while mean temperature conditions at bottom of the box was 17.3°C ± 6.6 *SD*). The heater was installed to mimic conditions in *E. nilssonii* roosts, which are often found in attics and roofing structured of heated, occupied houses, heated by direct sunlight (Rydell 1993b; Suominen et al. 2020). Before the breeding season of 2019 we installed a camera (D-link DSC-2670L) in the bat box, which provided motion-triggered video-recordings in addition to real-time observations (software mydlink, version 2.11.0, D-Link Corporation). Before the breeding season of 2020 we installed direction-sensitive infrared sensors (ChiroTec Tricorder 9006) at the box-entrance to record the timing of each emergence and return at night. Each pass registered by infrared sensors were checked against the motion-triggered video recordings to remove any false passes (i.e. if bats went to the exit, but did not fly out), to include passes that had been missed by infrared sensors, and to determine whether the recorded activity belonged to reproductive females or volant pups. Volant pups were distinguishable from adults on camera based on size and fur characteristics throughout their first flight week, after which the separation of volant pups and adults became increasingly challenging. Data on activity patterns therefore also include data from volant pups after they became undistinguishable from adults. However, pups adopt adult foraging patterns a week after gaining volancy (Fjelldal and van der Kooij 2024) and we therefore do not expect the inclusion of such individuals to cause large impacts in the data; nevertheless, this could be a source of variation to activity patterns observed towards the end of the breeding season.

The colony was monitored across the breeding period through daily box checks and video recordings to determine individual timing of arrival and parturition. When a newborn pup was detected, the mother was identified by her ring-placement, forearm-length or wing-patterns, depending on which females that were present in the box that year. Although the handling and daily box checks may be considered invasive for wild bats, the colony was habituated to human activity and -handling given the nature of its origin.

Hourly weather data, including temperature (°C), wind speed (m/s) and precipitation (mm rain), were downloaded for each breeding season from the closest meteorological station (SN4460; 11.7 km from study site). Timing of sunset and sunrise was obtained through the r-package suncalc (Thieurmel and Elmarhraoui 2022), which was used to calculate night length (hours).

All methods were carried out in accordance with relevant guidelines and regulations. All handling was carried out under license from the Norwegian Environment Agency (2023/6818 and preceding project licenses). Ringing of bats was permitted under license 06-3742 from the Norwegian Environment Agency. Rehabilitation of abandoned bat pups was granted under license 2001/450 from the Directorate for Nature Management.

### Body mass throughout reproductive period

Biometric measures (weights to nearest 0.1 g using a digital mini-weight and forearm-length to nearest 0.1 mm using dial callipers) of reproductive females were generally taken in the morning throughout the breeding season (five to 15 times per individual per season). Measured females were offered few mealworms (∼1 gram per bat) after each measurement as compensation for the disturbance. We did not expect these supplementary feedings to strongly impact activity patterns on the following night given that mealworms were provided in the morning. Because the parturition dates were known for each female (see Study system), we calculated the time relative to parturition for each measurement. Body mass data were collected each year from 2017 to 2023.

All statistical analyses were performed in R (version 4.3.1). To account for non-independence between datapoints (several observations per female across years) we fitted a linear mixed model (LMM) using the lmer-function from the lme4 package (Bates et al. 2015) with female ID and year included as random effects. Female body mass was tested as the response against the fixed effects reproductive stage (‘gestation’ versus ‘lactation’), time relative to parturition (negative values indicated days left until parturition while positive values indicated days since parturition) and total rainfall from mid-May to mid-July (annual value). Temperature conditions for mid-May to mid-July had a correlation coefficient of -0.95 with total rainfall and could therefore not be tested in the same model. Time relative to parturition was tested in interaction with reproductive stage. We also tested for the presence of non-linear temporal effects by including a quadratic effect of time relative to parturition in interaction with reproductive stage. We compared Akaike’s information criterion corrected for small sample sizes (AICc; Burnham and Anderson 2003) of the model with and without the quadratic time effect to determine the best fit for the data; the AICc value improved by 39.3 when including the quadratic effect and it was therefore kept in the final model.

### Number of trips

The colony-level number of foraging trips per bat per night was calculated by dividing the total number of emergences or returns (using the highest value in cases where they differed, as some passes were not detected) per night logged by the infrared sensors on the number of adult bats.

We fitted a linear model with number of trips per bat per night as the response variable (we did not include year as a random effect as these data were only collected for three years, from 2021 to 2023). Explanatory variables included reproductive stage (‘gestation’ versus ‘lactation’, which here was based on the median parturition date for the colony per year), time relative to the median parturition date, night length, nightly mean temperature, nightly mean wind speed, and nightly total rainfall. Time relative to parturition and nightly temperature conditions were tested in interaction with reproductive stage. We also tested for the presence of non-linear temporal effects; the quadratic effect of time relative to median parturition date improved the AICc value of the model by 7.9 and was therefore kept in the final model.

### Duration of ‘main’ trips

We used the data on emergence and return activity logged by the infrared sensors (checked by the motion-triggered video recordings) as a proxy for the duration of ‘main’ foraging trips (first trip of the night), although *E. nilssonii* may allocate some time outside the roost on other activities (de Jong 1994). One of the ringed individuals was present in the colony every year of this data collection (2021-2023) and could be specifically identified on the video recordings. For unringed females, ranging from three to four each summer, we could not determine the ID of each bat leaving and returning; however, because the females all generally left the box in rapid succession in the evening we used the time from the first unringed bat left the box to the first unringed bat returned as a measure of ‘main’ trip duration, and similarly with the following bats emerging and returning, unless an unringed bat left the box again before they had all returned. For these analyses we only included observations from nights where unringed females emerged around the same time (time from first to last female < 16 minutes). This will have caused some uncertainty in the exact calculation of trip durations, but we deemed the data precise enough.

The duration of ‘main’ trips was fitted as the response variable in an LMM, with fixed effects including reproductive stage, time relative to the median parturition date, night length, nightly mean temperature, nightly mean wind speed, and nightly total rainfall. Time relative to median parturition date and nightly temperature conditions were included as interaction effects with reproductive stage. We tested for non-linear temporal effects and found that the quadratic time effect significantly improved the fit of the model (ΔAICc = 27.9). Date was included as a random effect in each model.

### Timing of first emergences

The first emergences per evening were recorded using the data collected by the infrared sensors, where we included all first emergences (number of emergences = number of adult bats in the box) or, in cases where the first bats returned swiftly after leaving the box, the number of emergences before a return was recorded. Timing of emergences were calculated as the number of minutes from sunset to emergence. Data were collected across four summers (2020-2023).

We fitted an LMM with the timing of emergences each night tested as the response variable. Fixed effects included reproductive stage, time relative to the median parturition date, night length, and mean temperature, mean wind speed and total precipitation during the last hour before sunset. We tested reproductive stage in interaction with time relative to the median parturition date and temperature. A test for non-linear temporal effects revealed that the model which included the quadratic time-effect was a better fit (ΔAICc = 8.9) than the model without the quadratic term included. Date was included as a random effect.

### Timing of last returns

Data on the last nightly colony-level returns to the box were collected through the infrared recordings, where we included all final returns (number of returns = number of adult bats in the box), or the number of returns after the last recording of a bat leaving the box. Timing of returns were calculated as the time left until sunrise when bats returned to the box. Data were collected across four summers (2020-2023).

We fitted an LMM with the timing of returns each night tested as the response variable. Reproductive stage, time relative to the median parturition date, night length, nightly mean temperature, nightly mean wind speed and nightly total rainfall were included as fixed effects. We tested time relative to the median parturition date and nightly temperature in interaction with reproductive stage. Our test for non-linear temporal effects revealed that the quadratic time-effect significantly improved the model fit (ΔAICc = 17.2).

### Data from one identified individual

One of the two ringed females in the colony was present across all breeding seasons and was easily identified on all camera-observations and box-checks given the ringed right wing of this individual (the second female was ringed on the left wing). To verify the temporal patterns at the colony level, we performed the same analyses as described in the previous Methods-sections on data from this identifiable individual, “Nikki”. This allowed us to look at the activity throughout the reproductive season at a finer scale than using median parturition dates based on all the females.

Additionally, as a measure of total foraging time, we calculated the proportion “Nikki” spent away from the roost each night (total time spent out divided by the night length), given the detailed information we had on each emergence and return of this individual. We fitted a linear model with proportion spent away as the response variable and reproductive stage, time relative to parturition date, night length, nightly mean temperature, nightly mean wind speed and nightly total rainfall as fixed effects. Time relative to the parturition date and nightly temperature were tested in interaction with reproductive stage. Adding time relative to parturition as a quadratic time-effect significantly improved the model fit (ΔAICc = 3.1).

## Results

### Body mass throughout reproductive period

Female body mass varied from 6.0 grams to 15.2 grams (mean: 10.8 grams ± 2.0 *SD*, N_obs_ = 276) throughout the study period and was dependent on reproductive stage (gestation versus lactation) and time of measurement relative to individual parturition date (Table 1a & Fig. 2a; individual body mass curves are shown in Fig. S1 in the Supplementary Materials). Pregnant females gained weight exponentially during the gestation period, from a predicted weight of 9.0 grams (95% CI [8.3, 9.7]) 40 days prior to giving birth, to a predicted weight of 14.4 grams (95% CI [13.8, 15.0]) at the day of parturition (predictions are made from model results; Fig. 2a). After giving birth, the body mass of lactating females decreased until they were observed to regain body mass again in the period between ∼40 days to ∼90 days after parturition (Fig. 2a), corresponding to the time from beginning of August to mid-September. Each year the colony dissolved by the end of July, but some of the females still occasionally revisited the bat box after this point in time.

**Table 1:** Model results from testing effects on the variation in **a)** body mass, **b)** number of foraging trips, and **c)** duration of main foraging trips in reproductive female E. nilssonii. The effect sizes for all lactation stage variables in each model are relative to effects of the gestation stage variables, with the p-value indicating whether the effects from the lactation stage are significantly different from the gestation stage.

| Variable | Estimate (SE) | t value | p value |
| --- | --- | --- | --- |
| **a) Model: Body mass throughout reproductive period** |  |  |  |
| Female ID (random effect) | 0.3 ($\pm 0.5$) | | |
| Year (random effect) | 0.08 ($\pm 0.3$) | | |
| Residual (random effect) | 1.3 ($\pm 1.1$) | | |
| Intercept (gestation stage) | 14.51 ($\pm 0.50$) | 29.0 | < 0.001 *** |
| Days relative to median parturition day | 0.24 ($\pm 0.02$) | 10.1 | < 0.001 *** |
| Days relative<sup>2</sup> | 0.003 ($\pm 0.0006$) | 4.4 | < 0.001 *** |
| Total rainfall | -0.008 ($\pm 0.003$) | -0.3 | 0.7783 |
| Lactation stage | -3.14 ($\pm 0.43$) | -7.2 | < 0.001 *** |
| Lactation stage × Days relative | -0.33 ($\pm 0.03$) | -10.1 | < 0.001 *** |
| Lactation stage × Days relative<sup>2</sup> | -0.001 ($\pm 0.0007$) | -1.9 | 0.0636 |
| **b) Model: Number of foraging trips** |  |  |  |
| Intercept (gestation stage) | 2.03 ($\pm 1.34$) | 1.5 | 0.1317 |
| Days relative to median parturition day | 0.07 ($\pm 0.02$) | 3.3 | 0.0012 ** |
| Days relative<sup>2</sup> | 0.002 ($\pm 0.0006$) | 3.5 | < 0.001 *** |
| Nightly Ta (gestation stage) | 0.11 ($\pm 0.03$) | 3.8 | < 0.001 *** |
| Night length | -0.18 ($\pm 0.23$) | | -0.79 0.4291 |
| Nightly wind | 0.13 ($\pm 0.07$) | 1.8 | 0.0745 |
| Nightly rain | 0.11 ($\pm 0.02$) | 5.2 | < 0.001 *** |
| Lactation stage | 0.86 ($\pm 0.48$) | 1.8 | 0.0802 |
| Lactation stage × Days relative | -0.13 ($\pm 0.03$) | -4.2 | < 0.001 *** |
| Lactation stage × Days relative<sup>2</sup> | -0.001 ($\pm 0.0008$) | -1.4 | 0.1747 |
| Lactation stage × Nightly Ta | -0.03 ($\pm 0.04$) | -1.0 | 0.3338 |
| **c) Model: Duration (minutes) of main foraging trip** |  |  |  |
| Date (random effect) | 794.4 ($\pm 28.2$) | | |
| Residual (random effect) | 1261.1 ($\pm 35.5$) | | |
| Intercept (gestation stage) | -147.24 ($\pm 94.86$) | -1.5 | 0.1283 |
| Days relative to median parturition day | -9.69 ($\pm 1.60$) | -6.0 | < 0.001 *** |
| Days relative<sup>2</sup> | -0.26 (0.04) | -5.9 | < 0.001 *** |
| Nightly Ta (gestation stage) | 8.12 ($\pm 1.98$) | 4.1 | < 0.001 *** |
| Night length | 25.89 ($\pm 16.43$) | | 1.6 0.1173 |
| Nightly wind | -6.44 ($\pm 5.09$) | -1.3 | 0.2081 |
| Nightly rain | -6.32 ($\pm 1.48$) | -4.3 | < 0.001 *** |
| Lactation stage | 86.62 ($\pm 33.85$) | 2.6 | 0.0116 * |
| Lactation stage × Days relative | 11.35 ($\pm 2.26$) | 5.0 | < 0.001 *** |
| Lactation stage × Days relative<sup>2</sup> | 0.29 ($\pm 0.06$) | 4.7 | < 0.001 *** |
| Lactation stage × Nightly Ta | -5.80 ($\pm 2.51$) | -3.2 | 0.0226 * |

**Figure 2:**
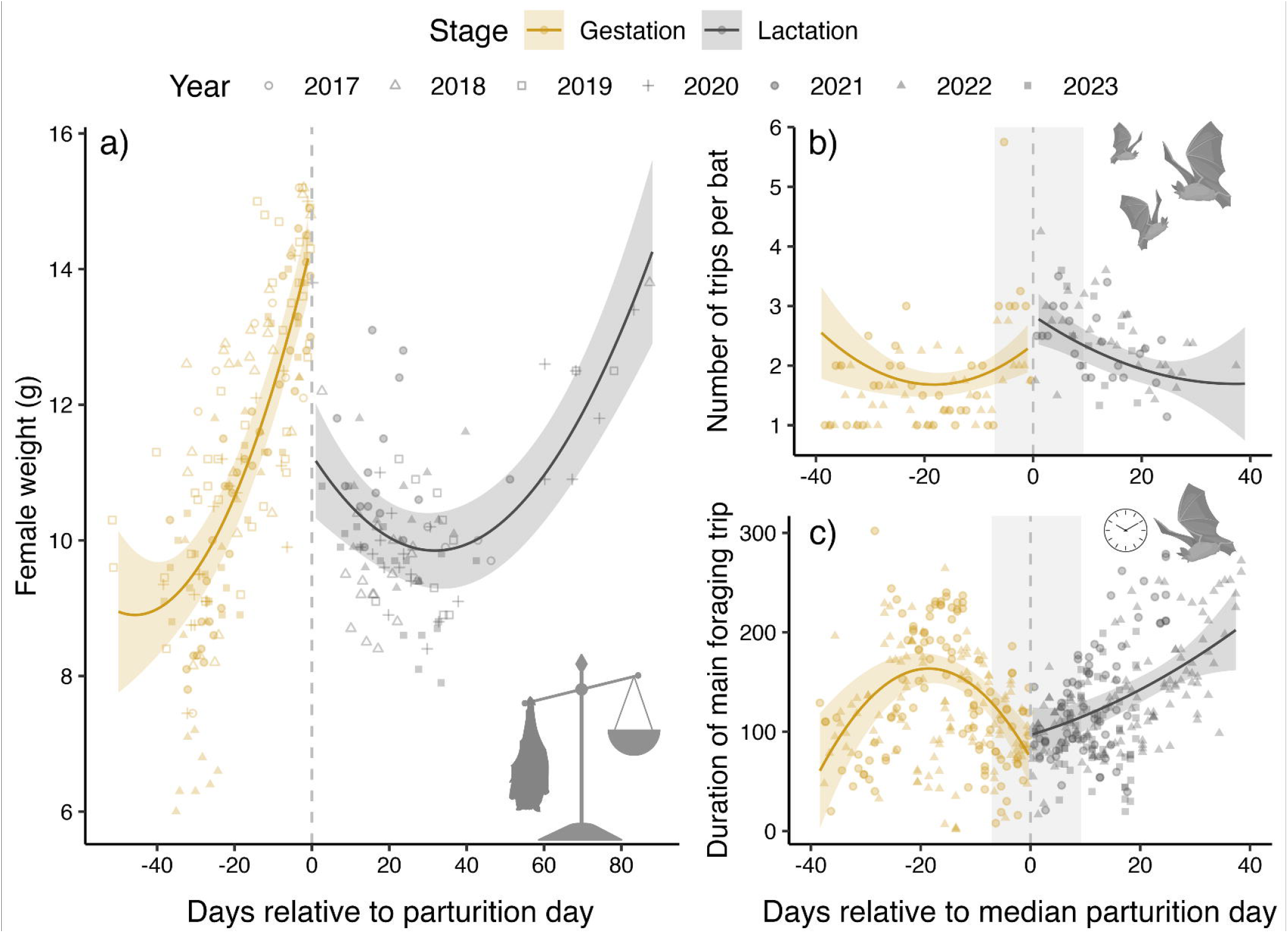
Temporal effects patterns during gestation (yellow) and lactation (grey), with regression lines and confidence intervals predicted from the models (Table 1) while datapoints show the raw data (different shapes indicate different years). Panes illustrate the variation in **a)** body mass in relation to the parturition date (for which the exact date per female was known), and variation in **b)** number of trips per bat and **c)** duration (minutes) of the main foraging trip in relation to the median parturition day (negative values indicate time left until the birth date). In panes b and c the shaded area indicates the time interval from the first pup born to the last pup born.

### Number of trips

The colony-level number of trips per bat (total number of trips/total number of bats) varied from 1 to 5.8 per night (mean: 2.0 trips ± 0.8 *SD*, N_obs_ = 148) throughout the study period and was dependent on reproductive stage, time relative to the median parturition date, nightly temperatures and nightly rainfall (Table 1b). Pregnant females generally had 1 to 2 trips per night, but the number of trips increased slightly during the last week before the median parturition date (Fig. 2b). At the time of the median parturition day, the number of trips was at its highest with a predicted number of 2.8 trips per night per lactating female (95% CI [2.4, 3.3]). After parturition, the number of trips declined steadily (Fig. 2b). Increasing nightly mean temperature and total nightly rainfall impacted the number of trips positively throughout the study period, resulting in more trips per bat on warm or rainy nights. We detected no difference in the temperature effect between reproductive stages.

### Duration of ‘main’ trips

The duration of ‘main’ foraging trips (first trip per bat each night) varied from 1.8 minutes to 5 hours (mean: 126.0 minutes ± 57.9 *SD*, N_obs_ = 405). The duration was strongly impacted by reproductive stage, time relative to the median parturition date, in addition to nightly mean temperatures and total nightly rain conditions (Table 1c). During the gestation period the duration of the main trip first increased up until around three weeks prior to the median parturition date. However, the duration of the main trip decreased after this point in time, reaching a predicted duration of 73.7 minutes (95% CI [45.3, 102.1]) at the time of the median parturition date (Fig. 2c). After parturition, the duration of the main trip increased rapidly again. Increased rainfall during the night led to a reduction in the duration of main trips, while warmer nights resulted in prolonged main trips during the gestation stage, but did not impact the duration of main foraging trips during the lactation stage (Table 1c).

### Timing of first emergences

Overall emergence time in the data varied from 17 minutes before sunset to 148 minutes after sunset (mean emergence: 32.3 minutes after sunset ± 19.7 *SD*, N_obs_ = 827) and was influenced by reproductive stage, time relative to median parturition, night length, and temperature before sunset (Table 2a). Females emerged significantly earlier during the lactation period than during the gestation period. The overall mean emergence time during the gestation stage (not accounting for any other effects) was 40.8 minutes after sunset (± 21.4 *SD*, N_obs_ = 339), while mean emergence time during the lactation stage was 26.4 minutes after sunset (± 16.0 *SD*, N_obs_ = 488). However, during the gestation period pregnant females emerged gradually earlier as they approached the date for median parturition, with a predicted emergence time of 43 minutes after sunset (95% CI [34.9, 51.3]) at the time of the median parturition date (Fig. 3a). During the lactation stage, females kept emerging gradually earlier until reaching the earliest predicted emergence time of 19.7 minutes after sunset (95% CI [15.3, 24.1]) 27 days after the median parturition date, after which they emerged increasingly later. Temperature conditions during the hour prior to sunset impacted pregnant and lactating females differently; while higher temperatures throughout the gestation period resulted in females emerging earlier from the roost, the opposite trend was observed during the lactating period, where increasing temperatures led to later emergences (Fig. 3b). Increasing night length by one hour resulted in bats delaying emergence by ∼14 minutes, accounting for other effects (Table 2).

**Table 2:** Model results from testing effects on the variation in **a)** timing of emergence and **b)** timing of return of reproductive female E. nilssonii. The effect sizes for all lactation stage variables in each model are relative to effects of the gestation stage variables, with the p-value indicating whether the effects from the lactation stage are significantly different from the gestation stage.

| Variable | Estimate (SE) | t value | p value |
| --- | --- | --- | --- |
| **a)** | **Model: Minutes from sunset to emergence** |  |  |
| Date (random effect) | 96.2 ($\pm 9.8$) | | |
| Residual (random effect) | 181.5 ($\pm 13.5$) | | |
| Intercept (gestation stage) | -25.83 ($\pm 27.74$) | -0.9 | 0.3527 |
| Days relative to median parturition day | 0.78 ($\pm 0.42$) | 1.9 | 0.0635 |
| Days relative<sup>2</sup> | 0.02 (0.01) | 1.9 | 0.0609 |
| Sunset Ta (gestation stage) | -0.81 ($\pm 0.44$) | -1.8 | 0.0706 |
| Night length | 14.07 ($\pm 4.85$) | 2.9 | 0.0041 *** |
| Sunset wind | 1.12 ($\pm 0.99$) | 1.1 | 0.2623 |
| Sunset rain | -1.25 (0.80) | -1.6 | 0.1210 |
| Lactation stage | -25.97 ($\pm 11.11$) | -2.3 | 0.0204 * |
| Lactation stage × Days relative | -2.73 ($\pm 0.60$) | -4.5 | < 0.001 *** |
| Lactation stage × Days relative<sup>2</sup> | 0.02 ($\pm 0.02$) | 1.0 | 0.2998 |
| Lactation stage × Sunset Ta | 2.12 ($\pm 0.65$) | 3.2 | 0.0014 ** |
| **b)** | **Model: Minutes from return to sunrise** |  |  |
| Date (random effect) | 1013.5 ($\pm 31.8$) | | |
| Residual (random effect) | 949.9 ($\pm 30.8$) | | |
| Intercept (gestation stage) | 238.59 ($\pm 81.03$) | 2.9 | 0.0036 ** |
| Days relative to median parturition day | 6.22 ($\pm 1.31$) | 4.8 | < 0.001 *** |
| Days relative<sup>2</sup> | 0.16 (0.03) | 4.7 | < 0.001 *** |
| Nightly Ta (gestation stage) | -15.26 ($\pm 1.58$) | -9.7 | < 0.001 *** |
| Night length | 7.79 ($\pm 14.27$) | 0.5 | 0.5857 |
| Nightly wind | 2.31 ($\pm 4.28$) | 0.5 | 0.5905 |
| Nightly rain | 1.43 ($\pm 1.28$) | 1.1 | 0.2658 |
| Lactation stage | -207.76 ($\pm 29.55$) | -7.0 | < 0.001 *** |
| Lactation stage × Days relative | -5.83 ($\pm 1.92$) | -3.0 | 0.0027 ** |
| Lactation stage × Days relative<sup>2</sup> | -0.17 ($\pm 0.05$) | -3.4 | < 0.001 *** |
| Lactation stage × Nightly Ta | 12.47 ($\pm 2.06$) | 6.1 | < 0.001 *** |

**Figure 3:**
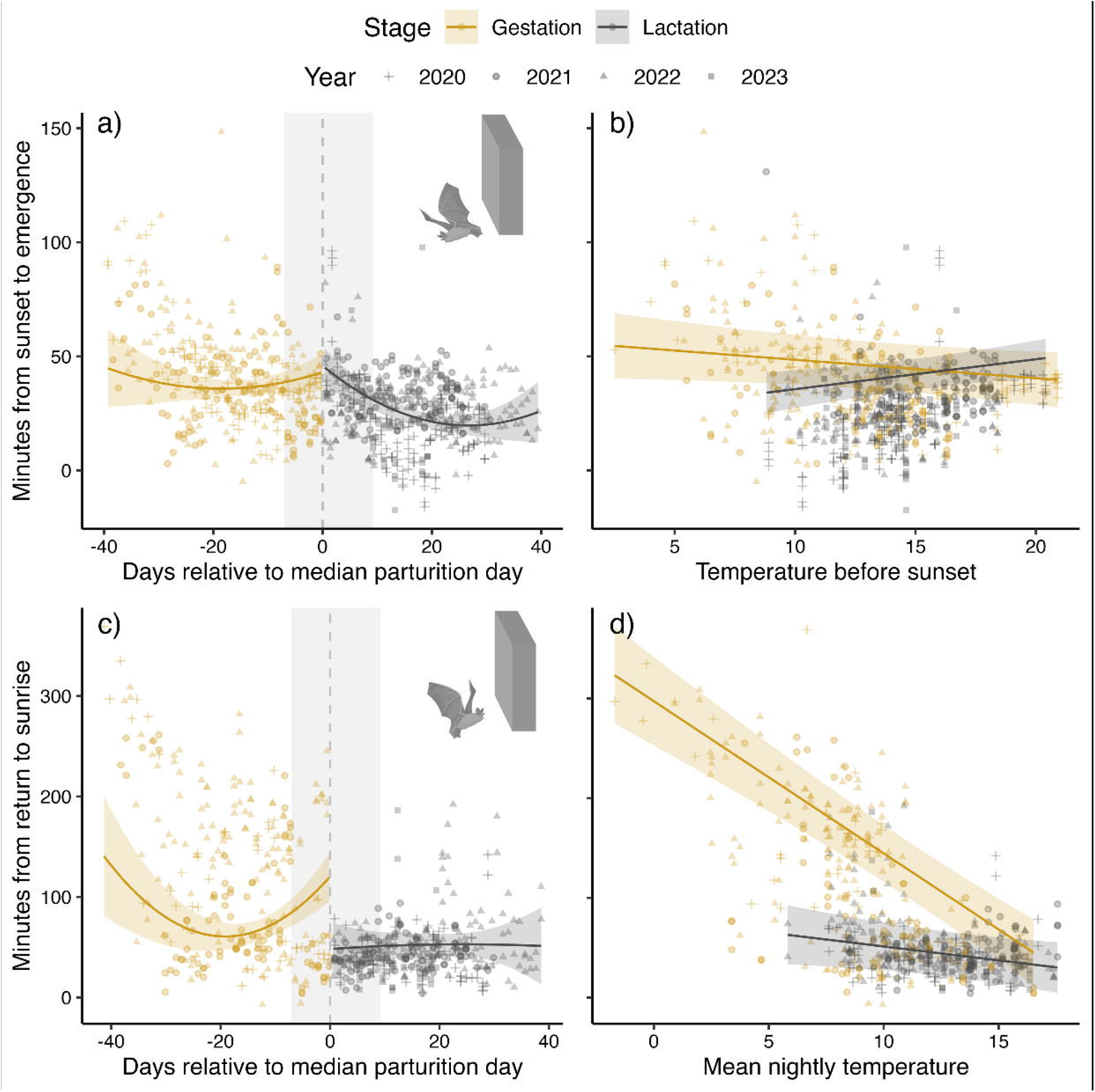
Temporal effects on emergence and return time during gestation (yellow) and lactation (grey), with regression lines and confidence intervals predicted from the models (Table 2) while datapoints show the raw data (different shapes indicate different years). **a)** Timing of emergence from the roost in relation to the median parturition day (negative values indicate time left until the birth date). Lower y-values indicate earlier emergences (0 marks sunset). **b)** Timing of emergence from the roost in relation to the mean temperature measured the hour before sunset. **c)** Timing of return to the roost in relation to the median parturition day. Lower y-values indicate later return (0 marks sunrise). **d)** Timing of return to the roost in relation to mean nightly temperatures. The shaded area in panes a and c indicates the time interval from the first pup born to the last pup born.

### Timing of last returns

Overall time of return to the roost varied from 369 minutes before sunrise to 7 minutes after sunrise (mean return time: 76.5 minutes before sunrise ± 65.8 *SD*, N_obs_ = 625). The timing of return to the roost was affected by reproductive stage, time relative to median parturition and mean nightly temperature (Table 2b). Lactating females generally returned later (i.e. closer to sunrise) than pregnant females. Overall mean return time during the gestation period (not accounting for other effects) was 115 minutes prior to sunrise (± 79.2 *SD*, N_obs_ = 272), while mean return time during lactation was 46.5 minutes before sunrise (± 28.0 *SD*, N_obs_ = 353). During the gestation period pregnant females returned gradually later to the roost as the time of median parturition approached, until reaching the latest predicted returns of 60.7 minutes before sunrise (95% CI [47.6, 73.9]) 19 days prior to parturition, at which point the females started returning gradually earlier again (Fig. 3c). After the median parturition date females were no longer affected by the time relative to parturition and returned considerably closer to sunrise. Increasing nightly temperatures generally resulted in females returning later; however, pregnant females were significantly more sensitive to changes in nightly temperatures than lactating females (Fig. 3d).

### Data from a single identified individual

All the analyses performed on data from the single identified individual “Nikki” confirmed the general patterns found at the colony level (Fig. S2 and S3 in the Supplementary Materials). We also confirmed that the data on this one identifiable bat did not drive the described activity patterns observed at the colony-level (see Fig. S5 in Supplementary Materials).

The actual time “Nikki” spent away from the box in total each night ranged from of 40 minutes to 4 hours during pregnancy (mean duration: 2.1 hours ± 0.94 *SD*, N_obs_ = 56) and from 46 minutes to 4.5 hours during lactation (mean duration: 3.1 hours ± 0.76 *SD*, N_obs_ = 80). When testing impacts on the proportion spent away from the roost at night, we found that the proportion varied between 0.11 to 0.78 (mean proportion: 0.49 ± 0.18 *SD*, N_obs_ = 136) and was affected by reproductive stage, days relative to parturition, mean nightly temperatures and total nightly rainfall (Table S1 in supplementary materials). The proportion spent away from the roost increased significantly from pregnancy to lactation (Fig. S4), with rainy nights decreasing this proportion, whereas warmer nights increased it. However, this positive effect of temperature was even stronger during pregnancy than during lactation (Table S1).

## Discussion

The results from our study show *E. nilssonii* at our monitored roost exhibit marked temporal differences in behaviour as a response to parturition, both in advance and after the event. Pregnant females show a decrease in time spent outside the roost during the three weeks leading up to parturition, correlating with body mass increase. Female bats both advance their exit from the roost as well as delay their return to the roost post parturition. These responses are dependent on temperature; however, lactating bats are less sensitive to temperature variability. They exit the roost earlier at low ambient temperature than prior to parturition and also extend duration outside the roost closer to sunrise at lower temperatures compared to bats prior to parturition. However, lactating females are prone to delaying roost exit at higher ambient temperatures. Females also increase the number of bouts outside the roost immediately after parturition, but the number of trips decreases and their length increases as time from parturition increases, likely because females can extend the duration of their trips between nursing as pups get older.

Lactation is an energetically demanding bodily function. Although we have no direct measurement of energy expenditure in the female bats studied here, we notice a continuous decline in female mass after parturition, with the predicted lowest mass occurring 32 days after giving birth. This decline in body mass is something that has not been reported in the literature previously, most likely due to the coarse nature of the data used in previous research (Kunz 1974; Mclean and Speakman 1999; Chaverri and Vonhof 2011). However, with lactation taking place for approximately 38 days in a closely related species, *E. fuscus* (Kunz 1971), and the costs of lactation being at their highest late on in the phase (Kurta et al. 1989), we can presume the costs of lactation to explain the decline in female body mass until pups begin shifting to arthropod diet. Although data on nursing were not systematically recorded in our study, we noted that females nursed their pups long after pups became volant. The three latest nursing-observations were made when pups were 25.4, 26.6 and 29.5 days old, which were, respectively, 11.1, 12.2 and 16.1 days after their first flight.

Previous research in *E. nilssonii* report lactating females usually making three foraging bouts each night, while pregnant and non-reproducing females made one or two (Rydell 1993a), with our results mirroring these findings. The additional trips can be attributed to the need to return to feed the pup at the roost during the night. However, with the pup growing rapidly and becoming volant already after two weeks (Fjelldal and van der Kooij 2024), the additional trips become less frequent with increasing pup age. Nevertheless, an additional seven days is still needed for the pups to adopt adult emergence and foraging patterns, which may reflect in some additional trips for the females as seen in our data set. The duration of individual trips also increases with pup age, which was also reported by Rydell previously (Rydell 1989).

Bats at northerly latitudes face a complication to their preparation for parturition with high enough nighttime temperatures for foraging activity only reached when nights are almost at their shortest close to summer solstice (Rydell 1989). With a need to avoid diurnal predators, this leaves the bats with only a limited time to forage during the night; total foraging time at night for one identifiable ringed female was observed to be ∼2 hours during gestation and ∼3 hours during lactation. The duration of trips outside the roost gradually decrease from three weeks prior to giving birth and is at its shortest around the time of parturition, an observation in the species also noted by Rydell (Rydell 1990). This coincides with the bats reaching maximum weight of up to 15 g in our dataset during late gestation, and the decrease in bouts of activity outside the roost can be attributed to bats avoiding the additional energetic costs of flying under heavy load and other factors related to late pregnancy. However, it is of note that parturition also appears to occur just as nighttime temperatures begin to reach their highest and fluctuation in temperature across nights is minimised (Fig. 3). This raises the notion that *E. nilssonii* females may be timing their parturition to take advantage of stable environmental conditions, with higher night time temperatures and increased arthropod abundance, at first instance to both cope with the elevated energetic demands of lactation as well as provide the longest possible time for pre-hibernation fattening before conditions deteriorate (Fjelldal et al. 2024).

We see a statistically significant shift to an earlier roost emergence during lactation, which is in accordance with previous studies (Swift 1980; Barclay 1989; Shiel and Fairley 1999; Russo et al. 2007). In addition, we find a delay in return to the roost prior to sunrise immediately in the wake of parturition. Based on results from prior research we can presume this behaviour to also reflect the increased energetic cost of lactation (Kurta et al. 1989; Duvergé et al. 2000). Although we cannot disclose the possibility of these bats also increasing their use of night time roosting outside the maternity roost, we can expect this behaviour to reflect an increase in foraging time to compensate for energetic needs of lactation.

As mentioned, the nature of our data does not allow us to take into account the possibility of bats resting outside the roost during the night: a behaviour which has, however, been documented in both pregnant and lactating *E. nilssonii* (Kosonen 2013). We acknowledge not being able to quantify the proportion of time used for foraging against time spent resting outside the maternity roost limit the validity of our findings. Another possible reason for extended foraging time post parturition would be the increase in night length following the coinciding summer solstice and parturition, but preliminary models taking night length into account denied this explanation (night length could not be tested in the final models due to collinearity with other variables). Furthermore, lactating females appear less sensitive to temperature, and they exit the roost early regardless of temperature, whereas pregnant bats delay their exit in colder conditions. This most likely again emphasizes the importance of energy intake for lactating females that must maintain elevated metabolism for milk production. In fact, it appears that at lower temperatures, gestating bats forage for shorter period in total per night, and can make use of torpor during inclement weather conditions (Willis et al. 2006). However, the responses to temperature in gestating and lactating females cease to differ at higher temperatures. Although our study is lacking in data on foraging activity and insect availability *per se*, existing literature allows us to deduct a plausible explanation for this maybe that potential energy intake at warmer ambient temperature is higher due to increased insect activity (Swift 1980; Rydell et al. 1996), and lactating females do not need to risk predation as a consequence of early roost exit. Similarly, at higher ambient temperatures, lactating females match their return to the roost with gestating females, avoiding late returns closer to sunrise, which can be attributed to females having had enough to eat during a warm night and not having to risk elevated predation associated with extended foraging and late return to roost. Indeed, any future studies on the topic would greatly benefit from parallel sampling of insect availability to validate our presumptions.

Data from the single ringed individual, “Nikki”, through which we could investigate the exact effects of time relative to parturition, confirmed the general activity patterns observed across the breeding season in this colony. However, the temporal effect on the number of trips made per night during gestation and lactation was markedly stronger at the individual level (see Fig. 1b versus Fig. S2b). This illustrates the immediate and substantial shift in activity the transition from gestation to lactation imposes on mammals. Such effects may be somewhat masked at the population-level in breeding colonies (Duvergé et al. 2000) due to the variation in individual timing of parturition (Sunga et al. 2023; Fjelldal and van der Kooij 2024). This can be difficult to account for in field studies, unless individuals are marked and closely monitored; data from the overlapping period when colonies or groups consist of both pregnant and lactating females should therefore be investigated with care. Furthermore, the plastic temporal responses during gestation and lactation found in our study highlights the importance of using finer scales than pre- and post-parturition, if possible.

Placing our results along a volume of research on *E. nilssonii* (Rydell 1992a; Rydell 1993a; Speakman et al. 2000; Suominen et al. 2020; Fjelldal et al. 2023; Suominen et al. 2023; Fjelldal et al. 2024; Fjelldal and van der Kooij 2024; Suominen et al. 2024) allows for a reserved interpretation of our findings in a larger context. Seasonal environments impose fluctuating selection on life history traits that can elicit adaptive responses (Varpe 2017). Although the summer is very short at northerly latitudes, the species has evolved behaviours and adaptations allowing it to breed at high latitudes. For instance, tolerance to light (Rydell 1992b; Frafjord 2021) and an ability to forage in a variety of environments (Vasko et al. 2020), including those more closed and shaded, may allow *E. nilssonii* to exploit the insects available with increasing nighttime temperatures earlier than other bat species. Early parturition maximises the time available for the pups to prepare for winter (Fjelldal et al. 2024; Suominen et al. 2024) and may ultimately be facilitated through the combination of multiple factors such as being able to take advantage of winter feeding opportunities (Blomberg et al. 2021) a generalist arthropod diet providing high foraging success (Vesterinen et al. 2018) in addition to the aforementioned tolerance to light (Speakman et al. 2000). Furthermore, the species exclusively utilize heated buildings as maternity roosts in Fennoscandia (Rydell 1993b; Suominen et al. 2023), and in our case a heated bat box, further enhancing the ability of the species to cope with the short summer seasons. The mechanisms behind these adaptations warrant further investigation, particularly in with respect to the impacts climate change may have on the phenology of the species and community it currently thrives in (Visser and Both 2005; Festa et al. 2023).

Finally, the results of our study have direct implications for conservation planning and management. Identifying roost sites and quantifying the number of bats roosting in them is an integral part of environmental impact assessments. The timing of these surveys can affect the results gained to a large degree and even result in previously identified occupied roosts being identified as abandoned, if activity at the roost is low, e.g. during time of parturition. Furthermore, our results highlight the expeditious nature of maternal care in bats at northerly latitudes, and in *E. nilssonii*, in particular. Maternity roost sites may only be in use for five to six weeks, which means the location and identification of these should be prioritized to the weeks preceding and following summer solstice.

## Author contributions

TML, MAF and JK designed the study, JK collected data, MAF curated and analysed the data, TML, MAF and JK interpreted the results, TML and MAF wrote the first draft of the manuscript, all authors participated in producing the final draft.

## Data availability statement

All data presented in this study, along with the r-codes used to analyse the data, are available through the following open access Zenodo-repository: https://github.com/tmlill/mammabats

## Additional Information

The authors have no competing interests.

## Acknowledgements

We sincerely thank Keith Redford and Karl Kugelschafter (Chirotec) for assisting with the infra-red based surveillance of the bat box, Ronny Steen (NMBU) for financing and installing the video surveillance, Rune Sørås and Anke Kirkeby and several volunteers for helping out with collecting data. We thank the Finnish Research Council and Jenny ja Antti Wihurin Säätiö for funding the research.

