## Supplementary material for "Behavioural changes in female northern bats (*Eptesicus nilssonii*) across gestation and lactation"

Supplementary Materials

**Individual body mass during reproduction**

Figure S1 illustrates the variation in body mass of individual females against date (a) and time relative to parturition (b) across the breeding seasons of 2017-2023.

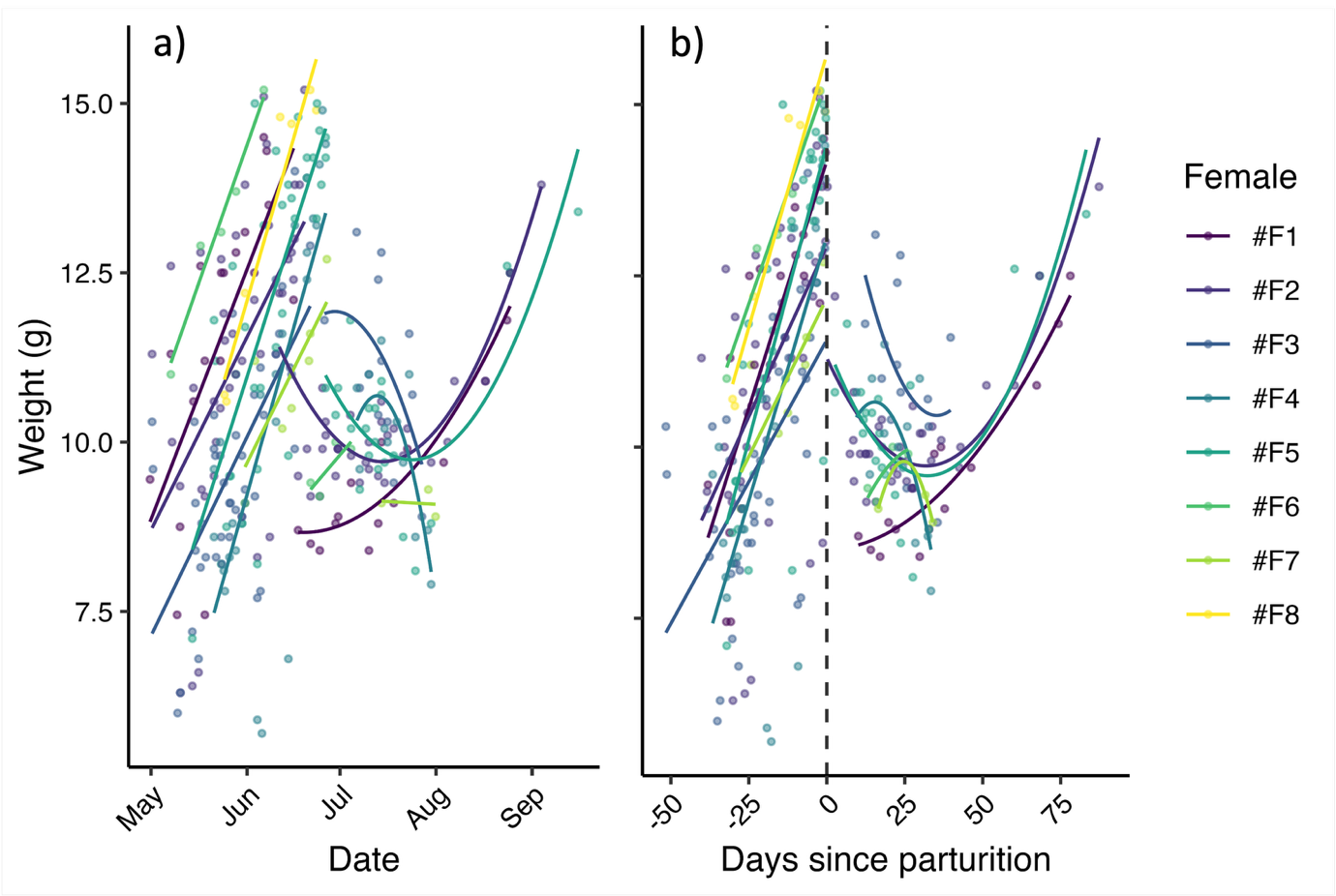

**Figure S1**: Body mass (g) of individual female *E. nilssonii* during the reproductive period (data from all years are included). **a)** Body mass shown against date, with the linear increase for each individual illustrating mass increase during gestation, while quadratic effects illustrate individual body mass changes during lactation. **b)** Body mass shown against days since parturition (negative values indicate time left until giving birth).

**Data from “Nikki”, an identified individual in the breeding colony**

To verify the patterns detected in this population of *E. nilssonii*, we analysed data from a single ringed female that was easily identified on all video-recordings. This allowed us to look at the activity throughout the reproductive season at a finer scale than when using median parturition dates based on the whole population. All analyses of data collected on “Nikki” confirmed the temporal patterns of trip numbers and -durations (Fig. S2) and the temporal and environmental patterns of emergence and returns (Fig. S3) that we detected in the overall colony data. Additionally, our analysis of effects on the proportion “Nikki” spent away from the roost at night (used as a proxy for foraging time) revealed that reproductive stage, time relative to parturition, nightly temperature and nightly rainfall had a significant impact (Table S1). “Nikki” spent a significantly larger proportion of the night foraging during the lactation phase compared to during gestation (Fig. S4). Increased total rainfall decreased the proportion spent away from the box at night, while nightly mean temperature had a positive effect; however, the sensitivity to variation in nightly temperature conditions was significantly higher during the gestation phase than during lactation (Table S1).

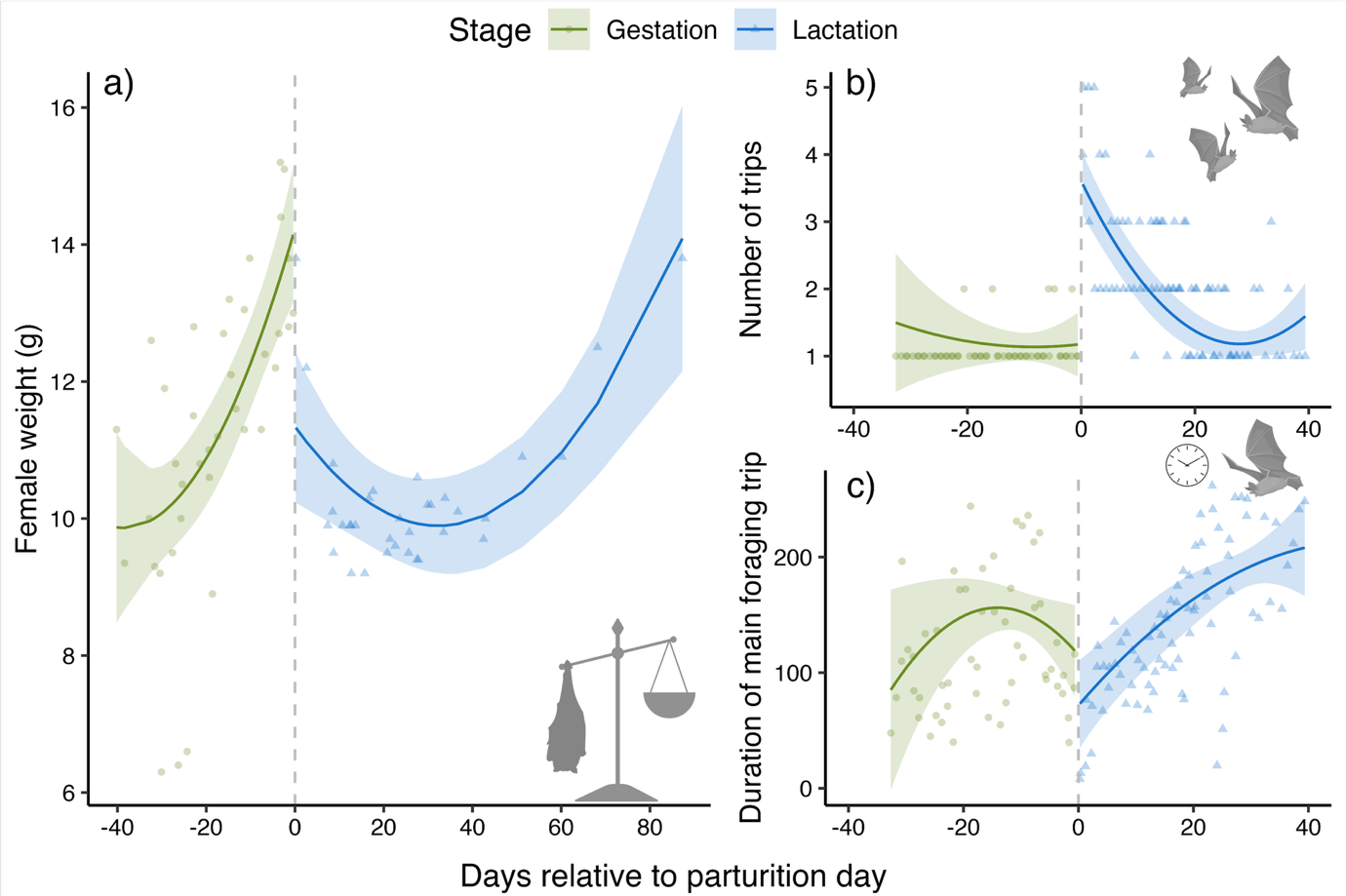

**Figure S2**: Temporal patterns during gestation (green) and lactation (blue) of a single identified individual, “Nikki”, in the colony. Panes illustrate the variation in **a)** body mass in relation to the parturition date, and variation in **b)** number of trips per night and **c)** duration (minutes) of the main foraging trip in relation to the parturition day (negative values indicate time left until the birth date).

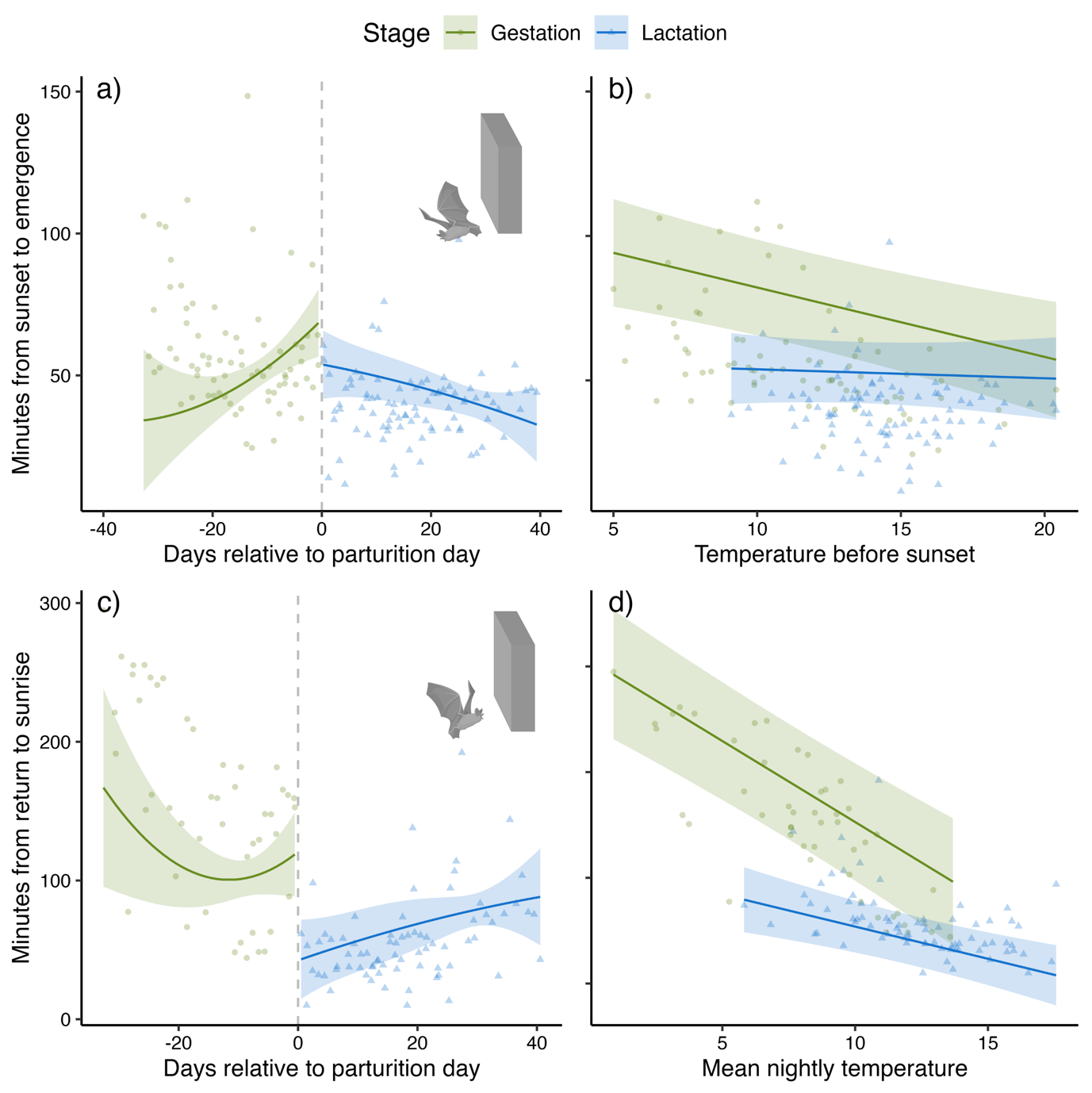

**Figure S3**: Temporal effects on emergence and return time during gestation (green) and lactation (blue) of a single known individual in the colony. **a)** Timing of emergence from the roost in relation to the parturition day (negative values indicate time left until the birth date). Lower y-values indicate earlier emergences (0 marks sunset). **b)** Timing of emergence from the roost in relation to the mean temperature measured the hour before sunset. **c)** Timing of return to the roost in relation to the parturition day. Lower y-values indicate later return (0 marks sunrise). **d)** Timing of return to the roost in relation to mean nightly temperatures.

**Table S1**: Model results from testing effects on the variation in the proportion spent away from the roost at night for the known individual, “Nikki”.

| Variable | Estimate (*SE*) | t value | *p* value |
| --- | --- | --- | --- |
| Intercept (gestation stage) | -0.13 (± 0.37) | -0.34 | 0.7327 |
| Days relative to parturition day | -0.02 (± 0.007) | -2.3 | 0.0257 * |
| Days relative^2 | -0.0005 (± 0.0002) | -2.2 | 0.0288 * |
| Nightly Ta (gestation stage) | 0.04 (± 0.007) | 5.9 | < 0.001 *** |
| Night length | 0.005 (± 0.06) | 0.1 | 0.9358 |
| Nightly wind | 0.017 (± 0.01) | 1.0 | 0.2973 |
| Nightly rain | -0.02 (± 0.005) | -3.3 | 0.0014 ** |
| Lactation stage | 0.33 (± 0.11) | 3.0 | 0.0030 ** |
| Lactation stage × nightly Ta | -0.02 (± 0.009) | -2.8 | 0.0054 ** |
| Lactation stage × Days relative | 0.025 (± 0.009) | 3.1 | 0.0023 ** |
| Lactation stage × Days relative^2 | 0.0003 (± 0.0002) | 1.4 | 0.1725 |

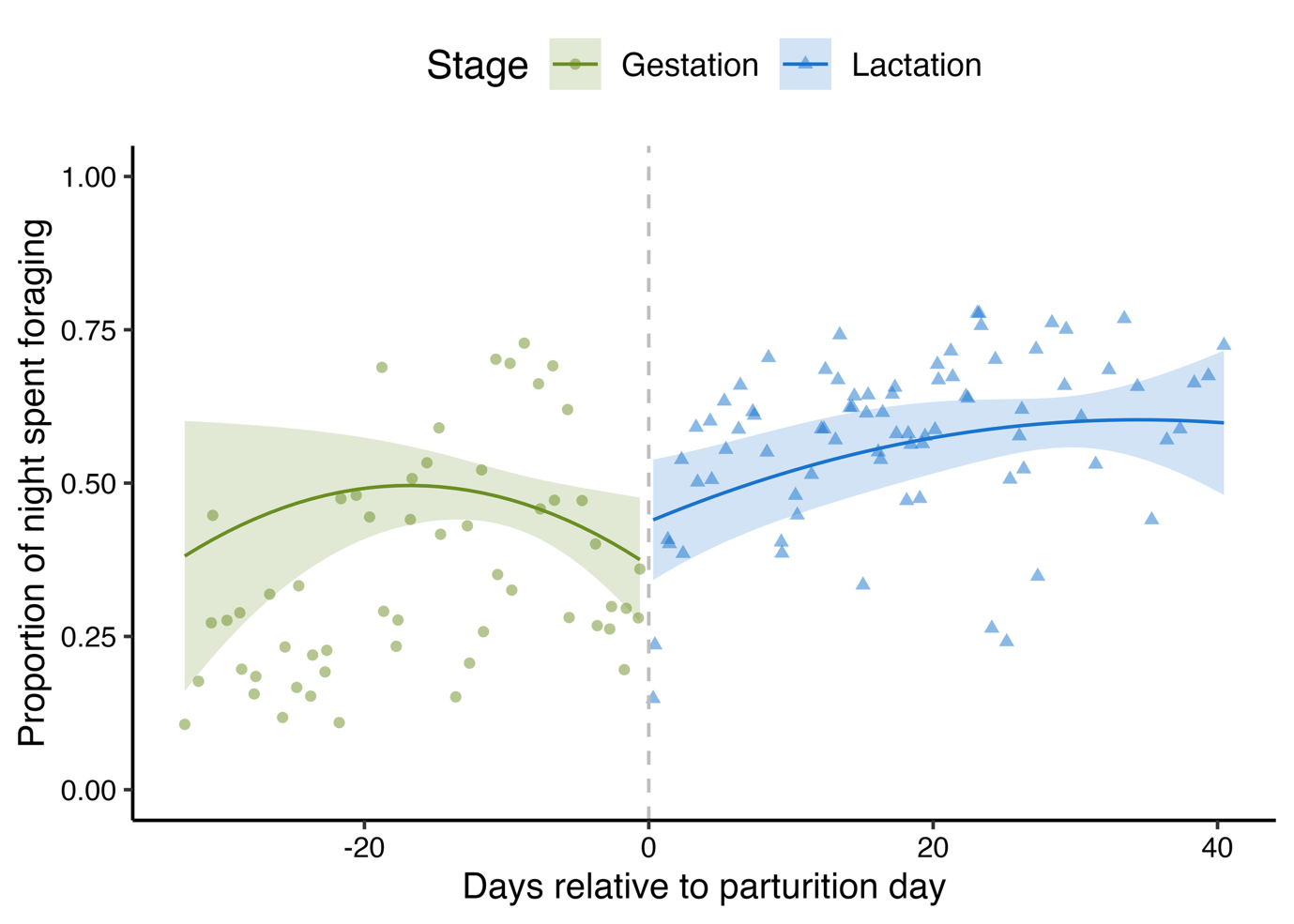

**Figure S4**: Variation in the proportion spent away from the roost at night as a response to time relative to parturition during gestation (green) and lactation (blue) of a single identified individual, “Nikki”, in the colony. Regression lines and confidence intervals are predicted from the model (Table S1) while datapoints show the raw data.
